# Systems level profiling of arginine starvation reveals MYC and ERK adaptive metabolic reprogramming

**DOI:** 10.1101/2020.01.13.904011

**Authors:** Caitlyn B. Brashears, Richa Rathore, Matthew Schultze, William R. Ehrhardt, Shin-Chen Tzeng, Brian A. Van Tine, Jason M. Held

**Affiliations:** Department of Medicine, Washington University in Saint Louis School of Medicine, St. Louis, MO 63110, USA; Siteman Cancer Center, Washington University in Saint Louis School of Medicine, St. Louis, MO 63110, USA; Department of Anesthesiology, Washington University in Saint Louis School of Medicine, St. Louis, MO 63110, USA

## Abstract

Arginine auxotrophy due to the silencing of argininosuccinate synthetase 1 (ASS1) occurs in many cancers, especially sarcomas. Arginine deiminase (ADI-PEG20) therapy exploits this metabolic vulnerability by depleting extracellular arginine, causing arginine starvation. ASS1-negative cells develop resistance to ADI-PEG20 through a metabolic adaptation that includes re-expressing ASS1. As arginine-based multiagent therapies are being developed, further characterization of the changes induced by arginine starvation is needed. In order to develop a systems-level understanding of these changes, activity-based proteomic profiling (ABPP) and phosphoproteomic profiling were performed before and after ADI-PEG20 treatment in ADI-PEG20-sensitive and resistant sarcoma cells. When integrated with previous metabolomic profiling (Kremer *et al*, 2017a), this multi-omic analysis reveals that cellular response to arginine starvation is mediated by adaptive ERK signaling, driving a Myc-Max transcriptional network. Concomitantly, these data elucidate proteomic changes that facilitate oxaloacetate production by enhancing glutamine and pyruvate anaplerosis, and altering lipid metabolism to recycle citrate for oxidative glutaminolysis. Based on the complexity of metabolic and cellular signaling interactions, these multi-omic approaches could provide valuable tools for evaluating response to metabolically targeted therapies.

## Introduction

The loss of a functional urea cycle in many types of cancer results from the silencing of argininosuccinate synthetase 1 (ASS1) expression (Kremer & Van Tine, 2017; Phillips *et al*, 2013; Khadeir *et al*, 2017; Keshet *et al*, 2018). This renders cancer cells dependent on extracellular arginine, as *de novo* arginine synthesis is reliant on ASS1 (Keshet *et al*, 2018). While the reason for this silencing has yet to be fully understood, current data suggest that it is beneficial for the production of biomass (Rabinovich *et al*, 2015; Cheng *et al*, 2018). Cancers that silence ASS1 have been shown to have a more aggressive clinical course, as silencing is associated with poorer overall survival and metastasis-free survival in numerous subtypes of cancer (Kobayashi *et al*, 2010; Huang *et al*, 2013; Qiu *et al*, 2014; Allen *et al*, 2014).

To exploit this metabolic deficiency, multiple arginine destruction enzymes have been developed, including: arginase, arginine decarboxylase, and arginine deiminase (Zam, 2017; Kremer & Van Tine, 2017). The most clinically relevant is PEGylated arginine deiminase (ADI-PEG20), which is currently in clinical trials. ADI-PEG20 converts extracellular arginine to citrulline, which cannot be metabolized into arginine in the absence of ASS1. Early development of ADI-PEG20 as a monoagent failed to demonstrate a survival advantage, likely due to the rapid re-expression of ASS1 in tumors (Abou-Alfa *et al*, 2018). As our understanding of drug development in this field has progressed, it has become clear that most agents will fail as monoagents due to the adaptability of tumor metabolism (Kremer *et al*, 2017b). By understanding the metabolic reprogramming that ASS1-negative tumors undergo as they re-express ASS1, additional vulnerabilities have been identified in sarcomas (Kobayashi *et al*, 2010; Prudner *et al*, 2019a; Kremer *et al*, 2017b; Huang *et al*, 2013).

Proteomic profiling can assess multiple potential aspects of protein regulation, such as protein abundance or protein post-translational modifications (Held *et al*, 2010; Rardin *et al*, 2013; Atsriku *et al*, 2009). Alternatively, activity-based proteomic profiling (ABPP) can evaluate changes in protein activity (Cravatt *et al*, 2008), kinase activity (Duncan *et al*, 2012), or ligand binding events (Luzarowski & Skirycz, 2019). Changes in protein expression or protein-protein interactions invariably contribute to ABPP as well (Wolfe *et al*, 2013; Piazza *et al*, 2018a; Veyel *et al*, 2018). Ultimately, ABPP integrates multiple informative proteomic parameters and provides a broad view of proteomic regulation. For example, ABPP can identify adaptive kinomic changes based on either altered kinase expression or activity (Duncan *et al*, 2012).

The mechanisms of developing resistance to arginine starvation in sarcomas have been partially defined, and include stabilization of nuclear cMyc (Prudner *et al*, 2019b), and increased glutamine anaplerosis in order to produce aspartate (Kremer *et al*, 2017a). In addition, others have examined mechanisms of ASS1 re-expression (Tsai *et al*, 2017; Long *et al*, 2017) and Deptor regulation (Ohshima *et al*, 2017). However, the underlying proteomic changes that initiate these events and coordinate metabolic reprogramming remain unknown. We pursued systems biology profiling to understand resistance to arginine starvation, as these approaches have proven effective in delineating the adaptive changes involved in highly pleiotropic phenotypes such as drug resistance (Zecena *et al*, 2018; Galluzzi *et al*, 2014), Myc activation, and various metabolic changes (Tomita & Kami, 2012; Schaub *et al*, 2018).

To understand ADI-PEG20-resistance of ASS1-negative sarcomas at a systems level, we performed multi-omic profiling using phosphoproteomics and activity-based proteomics, and coupled these data with existing metabolomic analyses (Kremer *et al*, 2017a). ADI-PEG20-senstive leiomyosarcoma cells (SKLMS1) have a much more dynamic phosphoproteomic response to ADI-PEG20 than a resistant angiosarcoma cell line (PCB-011). This includes increased phosphorylation of PDHA at Ser^293^, that inhibits entry of pyruvate into the mitochondrial TCA cycle via decarboxylation (Roche *et al*, 2004). ABPP profiling reveals that glutamine anaplerosis is facilitated by proteomic changes that drive the production of OAA by glutamine and by anaplerotic carboxylation of pyruvate, as well as the inhibition of lipid metabolism to recycle citrate to the TCA cycle. In addition, ABPP profiling reveals a Myc-Max transcriptional network that is regulated by adaptive changes in MAPK1 and MAPK2 upon ADI-PEG20 treatment in SKLMS1 cells. Therefore, we have demonstrated that multi-omic profiling can be utilized to delineate systems-level regulatory signaling networks mediating drug sensitivity and resistance. Due to the complex nature of metabolic and cell signaling interactions, these approaches could provide valuable tools for evaluating resistance and escape to metabolically targeted cancer therapies.

## Results

### Proliferative and morphologic changes of PCB-011 and SKLMS1 with ADI-PEG20 treatment

To systemically identify regulatory networks underlying resistance to ADI-PEG20 we examined two sarcoma cell lines, SKLMS1 (leiomyosarcoma) and PCB-011 (angiosarcoma). While SKMLS1 cells are sensitive to arginine starvation induced by ADI-PEG20, PCB-011 cells are rapidly resistant to ADI-PEG20 as indicated by their proliferative response over 72 hours (Fig. 1A and 1B). These data are consistent with the ASS1 expression in each cell line, as SKLMS1 is ASS1-negative, while PCB-011 is ASS1-positive (Fig. 1C), suggesting that this cell line can overcome extracellular arginine depletion. Finally, unlike PCB-011, there is a morphological change identified in the SKLMS1 cell line, as it becomes more spindle-like in response to arginine starvation (Fig. 1D). Cumulatively these data demonstrate the SKLMS1 is sensitive to arginine depletion with ADI-PEG20, while PCB-011 is not responsive.

**Figure 1.**
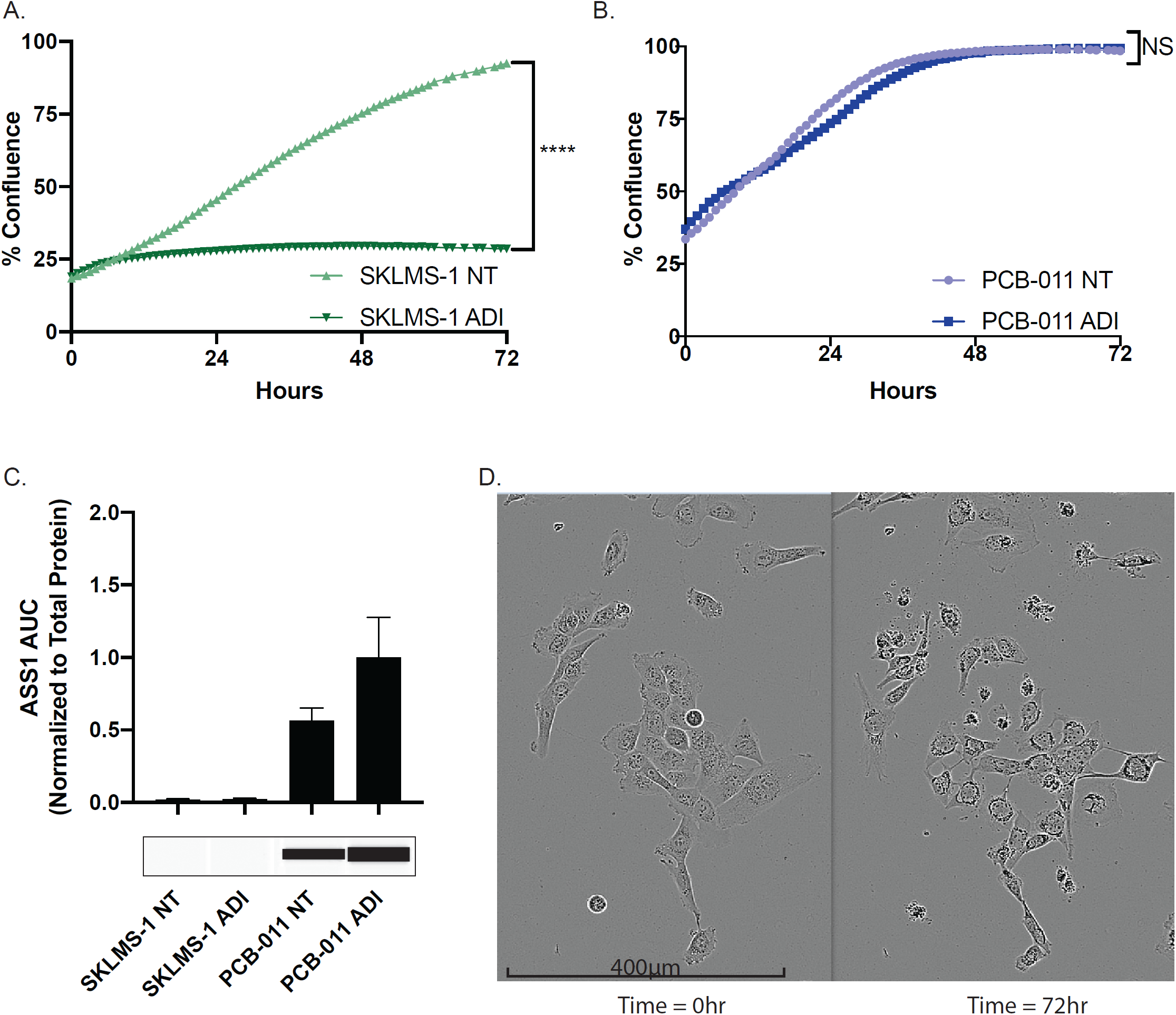
Proliferative and morphologic changes of PCB-011 and SKLMS1 with ADI-PEG20 treatment. **A)** *In vitro* cell proliferation response to extracellular arginine deprivation with ADI-PEG20 in SKLMS1. Cell proliferation was measured using cell confluence on the IncuCyte Live Cell Analysis System. (n=3) ****p<0.0001. **B)** *In vitro* cell proliferation response to extracellular arginine deprivation with ADI-PEG20 in PCB-011. Cell proliferation was measured using cell confluence on the IncuCyte Live Cell Analysis System. (n=3) ****p<0.0001. **C)** Protein expression of ASS1 compared in untreated and ADI-treated SKLMS1 and PCB-011 at 72 hr. Cell lysates were analyzed with Simple Protein Wes automated western system. Band density differences were plotted as ASS1 area under the curve normalized to total protein in the capillary (representative n=3); data are represented as mean + SD. **D)** Representative (n=3) DIC images of SKLMS1 at 0 hr and 72 hr of ADI-PEG20 treatment. Images were collected on the IncuCyte Live Cell Analysis System at 20x magnification.

### Phosphoprotemomic changes as a result of ADI-PEG20 treatment in sensitive and resistant cell lines

Glutamine and glucose metabolic tracing has previously shown that SKLMS1 cells increase anaplerotic oxidative glutaminolysis to produce aspartate from oxaloacetate in response to ADI-PEG20-induced arginine starvation (Kremer *et al*, 2017a). In addition, Western blots of candidate metabolic regulatory proteins have shown decreased phosphorylation of PKM2 Y^105^ and LDHA Y^10^, and increased phosphorylation of PDH1 S^300^ in response to ADI-PEG20 (Kremer *et al*, 2017a). In order to gain insight into how proteomic adaptations in SKLMS1 cells promote altered metabolism and cell signaling to survive arginine deprivation, we performed ABPP using an ATP-resin (Wolfe *et al*, 2013) as well as phosphoproteomic profiling of SKLMS1 and PCB-011 cells with and without ADI-PEG20 treatment for 72 hours.

We first examined how the phosphoproteome of each cell line responded to ADI-PEG20 treatment. 2551 phosphopeptides were detected with a 1% FDR (ProteomeXchange repository identifier PXD017043). Notably, the phosphoproteomic response of ADI-PEG20-sensitive SKLMS1 cells was much more dynamic than PCB-011 cells, with more phosphopeptides up- and down-regulated by ADI-PEG20 (Fig. 2A). A heatmap of phosphopeptides altered by ADI-PEG20 treatment also indicates that the SKLMS1 phosphoproteome is more dynamic than ADI-PEG20-resistant PCB-011 cells when starved of arginine (Fig. 2B).

**Figure 2.**
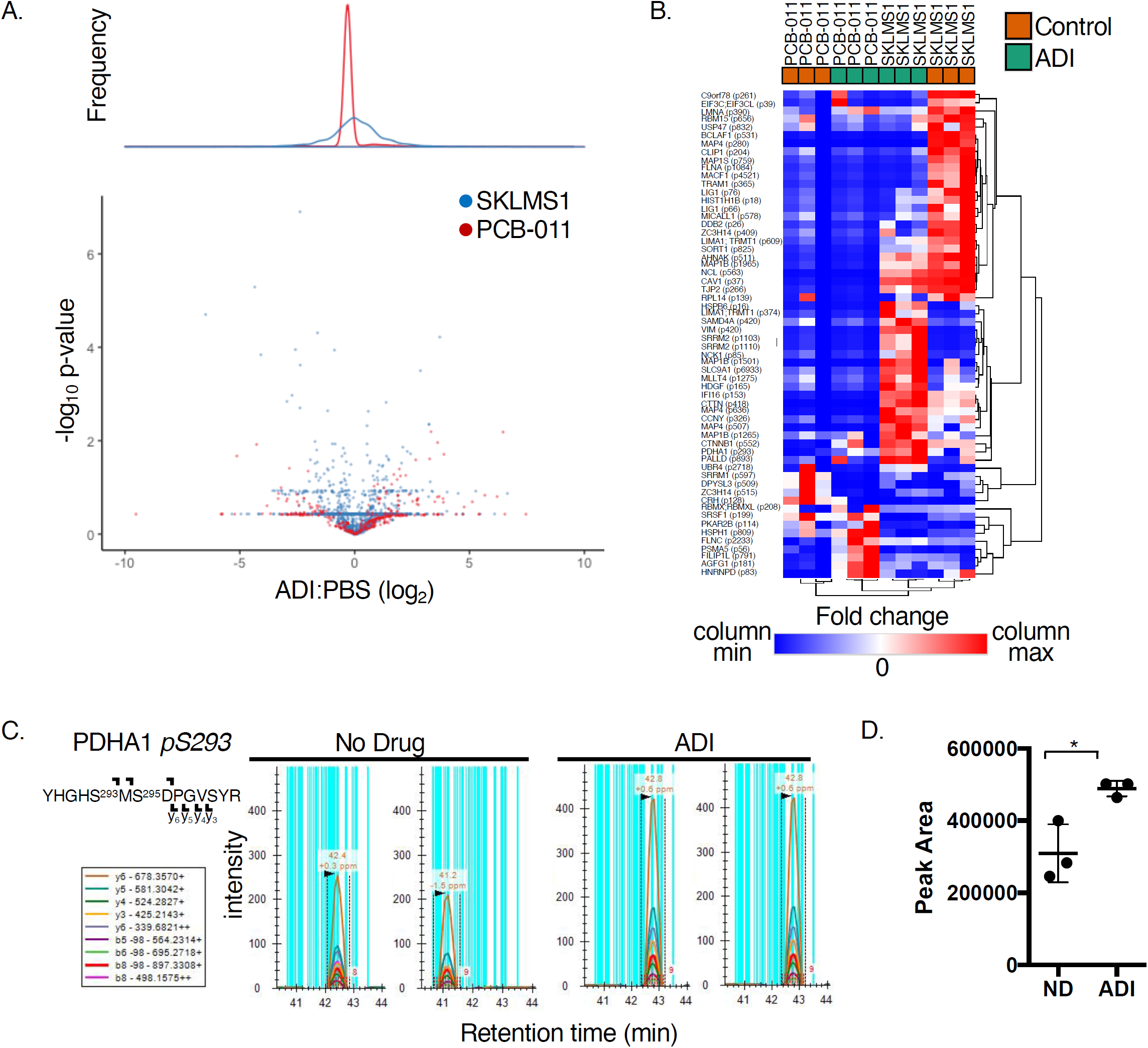
ADI-resistant SKLMS1 cells have a dynamic phosphoproteomic response to arginine starvation that regulates pyruvate dehydrogenase and proteins involved in cell morphology and contacts. **A)** Phosphoproteomic profiling of SKLMS1 and PCB-011 sarcoma cells upon ADI-PEG20-PEG (ADI-PEG20) treatment (72 h). **B)** Heatmap of proteins with significantly altered phosphorylation upon ADI-PEG20 treatment. Colors are assigned according to the directionality of deviation from no change (red, up; blue, down). Three biological replicates were analyzed for each cell per conditions. **C)** DIA-MS results for phosphorylation of S29 in PDHA1. **D)** Quantification of S29 phosphorylation in PDHA1 upon ADI treatment.

The phosphorylation of several notable proteins is uniquely upregulated in ADI-PEG20-resistant SKLMS1 cells. This includes phosphorylation of PDHA1 S^293^, which inhibits pyruvate entry into the TCA cycle through oxidative decarboxylation (Roche *et al*, 2004) and increases anaplerotic production of oxaloacetate (OAA) (Park *et al*, 2018). These data are consistent with previous metabolomic analysis (Kremer *et al*, 2017a). The increase in PDHA1 S^293^ phosphorylation was verified by DIA-MS (Held *et al*, 2013), which was able to clearly distinguish phosphorylation of S^293^ from other serines in the peptide (Fig. 2C). DIA-MS shows a significant (1.6 fold) increase in the expression of pS^293^ upon ADI-PEG20 treatment (Fig. 2D), paralleling the increase in PDHA S^300^ phosphorylation upon ADI-PEG20 treatment (Kremer *et al*, 2017a). Other phosphoproteins uniquely regulated by ADI-PEG20 treatment in ADI-PEG20-sensitive SKLMS1 cells include β-catenin, as well as the cell morphology and contact proteins LIMA1, VIM, MAP1B, MLLT4, CTTN, and PALLD, which is consistent with the change to a more fusiform morphology, as noted in ADI-treated SKLMS1 cells (Fig. 1D).

### SKLMS1 cells upregulate MAPK signaling and TCA proteins in response to ADI-PEG20, but downregulate lipid metabolism

ABPP detected 1,912 proteins (ProteomeXchange repository PXD017043), and substantial activity-based proteomic changes in both cell lines upon ADI-PEG20 treatment (Fig. 3A). A clustered heatmap including all proteins that were differentially regulated (p-value ≤0.05) reveals four distinct clusters, indicating that the proteomic response to ADI-PEG20 in each cell line is highly individualized (Fig. 3B).

**Figure 3.**
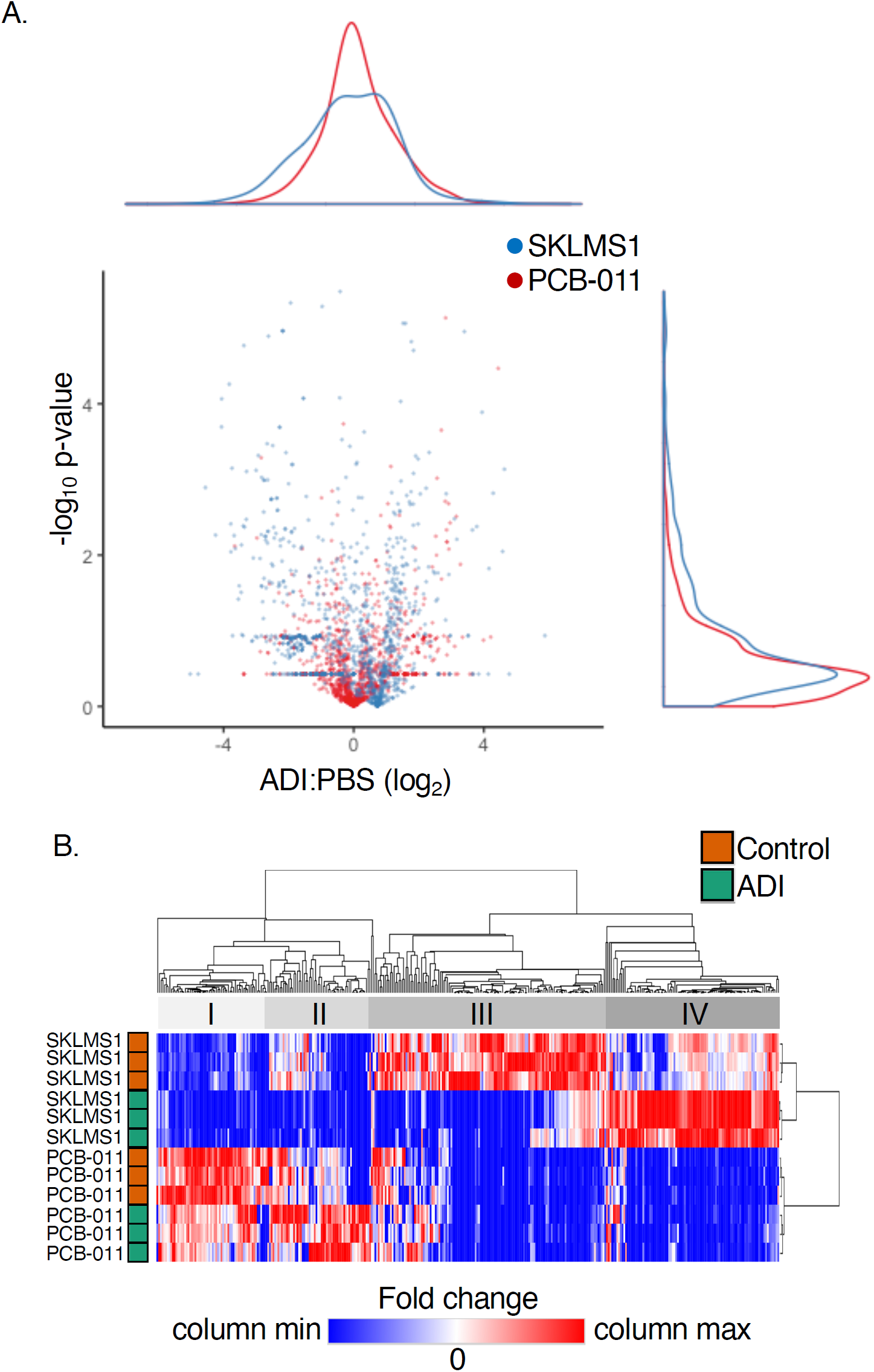
Activity based proteomic profiling reveals regulation of metabolism and kinases that are unique to ADI-resistant SKLMS1 cells upon arginine starvation. **A)** ABPP profiling of SKLMS1 and PCB-011 sarcoma cells upon ADI-PEG20-PEG (ADI-PEG20) treatment (72 h). **B)** Heatmap of proteins with significantly altered phosphorylation upon ADI-PEG20 treatment. Color choices are assigned according to the directionality of deviation from no change (red, up; blue, down). Three biological replicates were analyzed for each cell per conditions.

In order to determine the biological pathways and functions of proteins regulated by ADI-PEG20 treatment, we performed a gene set enrichment analysis (GSEA (Subramanian *et al*, 2005)). GSEA utilizes the fold changes of all proteins detected by ABPP, a more appropriate approach than filtering data by an arbitrary cutoff (Held, 2019). MAPK pathway and TCA cycle annotations were enriched in SKLMS1 cells after ADI-PEG20 treatment, but not in PCB-011 cells (Fig. 4A). Additionally, fatty acid, triacyglycerol, and ketone body metabolism was negatively regulated in SKLMS1, but not in PCB-011 cells (Fig. 4A). PCB-011 cells did not show strong positive enrichment of pathways, but showed negative regulation of translation and SRP-dependent co-translational protein targeting to membranes, which were also observed, albeit less strongly, in SKLMS1 cells (Fig. 4A). Enrichment plots and heatmaps of expression in SKLMS1 cells demonstrate coordinated regulation of proteins within the MAPK, TCA cycle, and fatty acid, triacylglycerol, and ketone body metabolism pathways upon ADI-PEG20 treatment (Fig. 4B). Regulation of the TCA cycle has been observed in SKLMS1 cells upon ADI-PEG20 treatment (Kremer *et al*, 2017a). Additionally, regulation of MAPKs and lipid metabolism has been observed in ADI-PEG20 sensitive melanoma (Long *et al*, 2013; Tsai *et al*, 2012). These results demonstrate that ADI-PEG20-sensitive SKLMS1 cells undergo metabolic and kinomic adaptation upon arginine starvation that is consistent with the existing literature.

**Figure 4.**
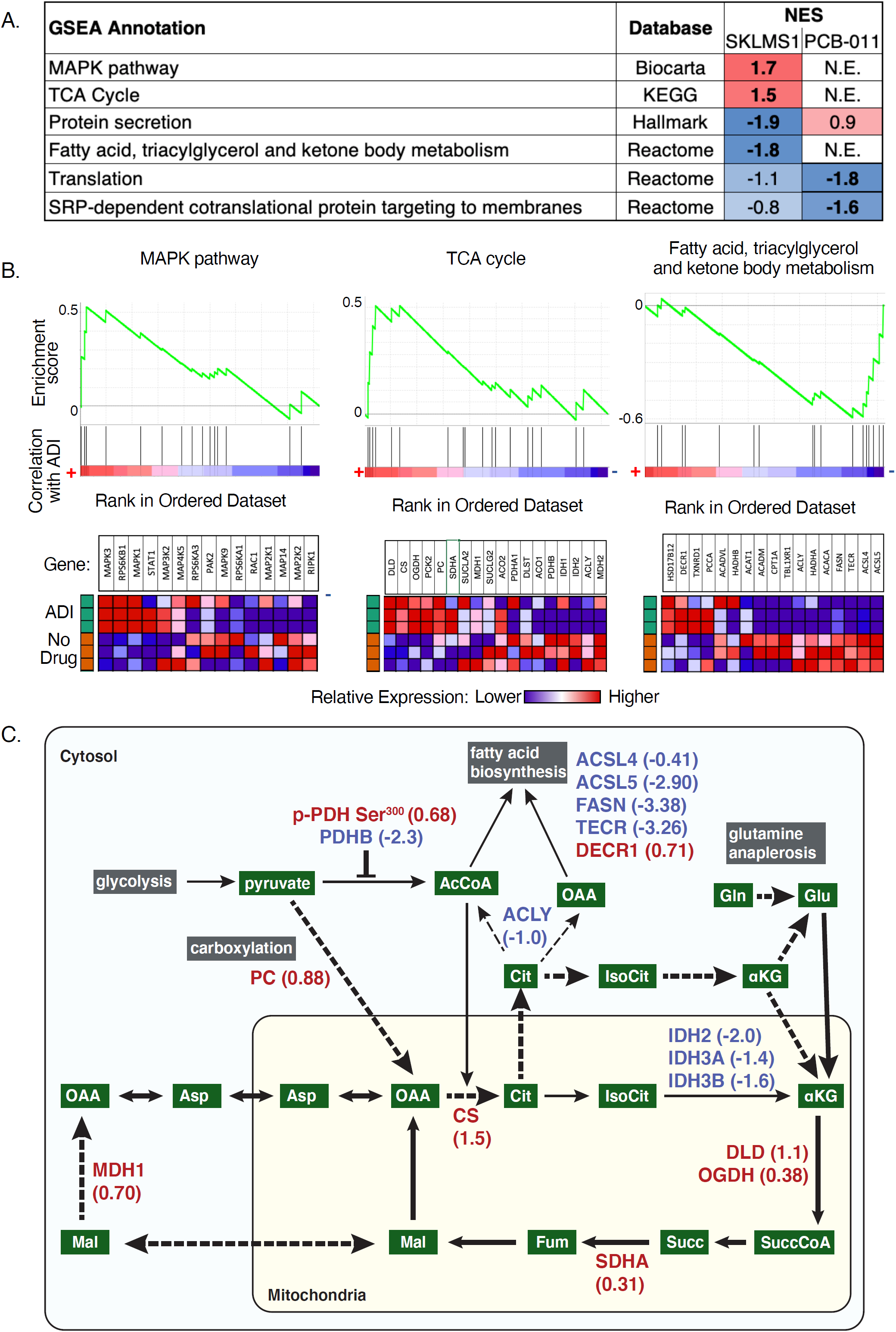
Arginine starvation induces MAPK signaling and coordinated proteomic alterations that promote glutamine anapleurosis, oxaloacetate formation, and inhibit lipid metabolism in SKLMS1 cells. **A)** Normalized enrichment scores (NES) of Annotations enriched from gene set enrichment analysis (GSEA) analysis for ABPP profiling of SKLMS1 and PCB-011 cells in response to ADI-PEG20 treatment. N.E. indicates Not Enriched and bold font indicates a false discovery rate <25%. **B)** Relative expression of individual genes in Annotations enriched in SKLMS1 cells. **C)** Schema of glycolysis, TCA cycle and lipid metabolism with ABPP and phosphoproteomic results (log2 fold change of ADI-treated compared to untreated, colors based on directionality of deviation from no change (red, up; blue, down)) in SKLMS1 cells, overlayed with known metabolic changes in SKLMS1 cells upon ADI treatment ((Kremer *et al*, 2017b), thick solid lines). Dashed lines indicate altered metabolism proposed by ABPP results. Colors are assigned according to the directionality of deviation from no change (red, up; blue, down).

### Arginine starvation induces coordinated proteomic alterations that promote glutamine anaplerosis and oxaloacetate formation, and inhibit lipid metabolism in SKLMS1 cells

ADI-PEG20 treatment increases glutamine anapleurosis through the TCA cycle, forming oxaloacetate to produce aspartate (Kremer *et al*, 2017a). However, the specific proteomic alterations that facilitate this metabolic rewiring remain unknown. In order to provide a more complete understanding of the mechanisms underlying the adaptive metabolic rewiring in response to arginine starvation, we focused on the ABPP regulation of individual proteins in the differentially regulated TCA cycle and fatty acid, triacylglycerol and ketone body metabolism annotations in SKLMS1 cells upon arginine starvation (Fig. 4B).

Multiple proteomic alterations support enhanced glutamine anapleurosis and utilization of oxaloacetate (OAA). First, multiple enzymes that drive glutamine anapleurosis to OAA were upregulated, including DLD, OGDH, and SDHA (Fig. 4C). In addition, IDH2, IDH3A, and IDH3B are substantially downregulated, blocking the reverse activity of the TCA cycle that can occur in cancer (Filipp *et al*, 2012), and further directing αKG towards OAA. Second, PDHB levels are decreased and pyruvate carboxylase (PC) levels are increased. Together with the increased phosphorylation of PDH S^300^ (Fig. 2D), which inhibits PDH activity (Roche *et al*, 2004), these findings suggest that more pyruvate is directly converted to OAA via anaplerotic carboxylation upon arginine deprivation (Fig. 4C). Third, ABPP finds that while citrate synthase (CS) is upregulated, metabolism of citrate to fatty acids is likely blocked due to decreased ACLY, ACSL4, ACSL5, FASN, and TECER. Since conversion directly to mitochondrial αKG is blocked by substantially decreased IDH2, 3A, 3B levels (Fig. 4C), though cytoplasmic IDH1 is largely unchanged (ProteomeXchange repository identifier PXD017043), citrate is likely shunted cytoplasmically to αKG or glutamate and back into the TCA cycle to undergo another anaplerotic cycle. However, the exact route cannot be determined from these results. Fourth, while mitochondrial malate dehydrogenase (MDH2) is largely unchanged upon arginine starvation, MDH1 is upregulated by 1.6 fold, suggesting that OAA may be preferentially formed from malate that has been exported to the cytoplasm (Fig. 4C). Taken together, the results from the ABPP analysis suggest that glutamine-based production of OAA and aspartate is driven by three potential pathways: increased TCA cycle activity, anaplerotic carboxylation of pyruvate, and inhibition of lipid metabolism that recycles cytoplasmic citrate back to the TCA cycle.

### Arginine starvation induces adaptive kinomic changes driving MYC-MAX activation in SKLMS1 cells

The regulation of metabolic adaptation and reprogramming is highly pleiotropic, coordinating regulation of kinases, transcription factors, and other proteins across multiple signaling pathways (Brooks Robey *et al*, 2015). Kinases are key transducers of signaling pathways, often clinically actionable, and can be unbiasedly profiled by ABPP (Duncan *et al*, 2012; Xiao & Wang, 2016). Since GSEA indicated that MAPKs were uniquely upregulated in SKLMS1 cells upon arginine starvation (Fig. 4A and B), we further investigated the kinomic changes of SKLMS1 and PCB-011 cells in response to ADI-PEG20. Consistent with phosphoproteome regulation (Fig. 3A and B), ADI-PEG20-resistant SKLMS1 cells have a much more dynamic response to ADI-PEG20 treatment than PCB-011, and the patterns of kinases that are regulated is distinct (Fig. 5A). Kinases with significant changes in protein expression (p≤0.05) are shown in Fig. 5B which includes 14 kinases in SKLMS1 cells, compared to two in PCB-011 cells. Notably, while SKLMS1 cells do not harbor activating mutations in ERK or AKT/mTOR signaling, ERK1 and ERK2 (MAPK3 and MAPK1, respectively) along with the ERK substrate p70 S6 kinase (Cravatt *et al*, 2008) have the largest increases upon ADI-PEG20 treatment by ABPP profiling. Each of these kinases promotes tumor growth and is capable of reprograming cellular metabolism (Papa *et al*, 2019; Um *et al*, 2004). Significantly, ERK activation has been implicated in the escape mechanism to ADI-PEG20 in melanoma (Long *et al*, 2013).

**Figure 5.**
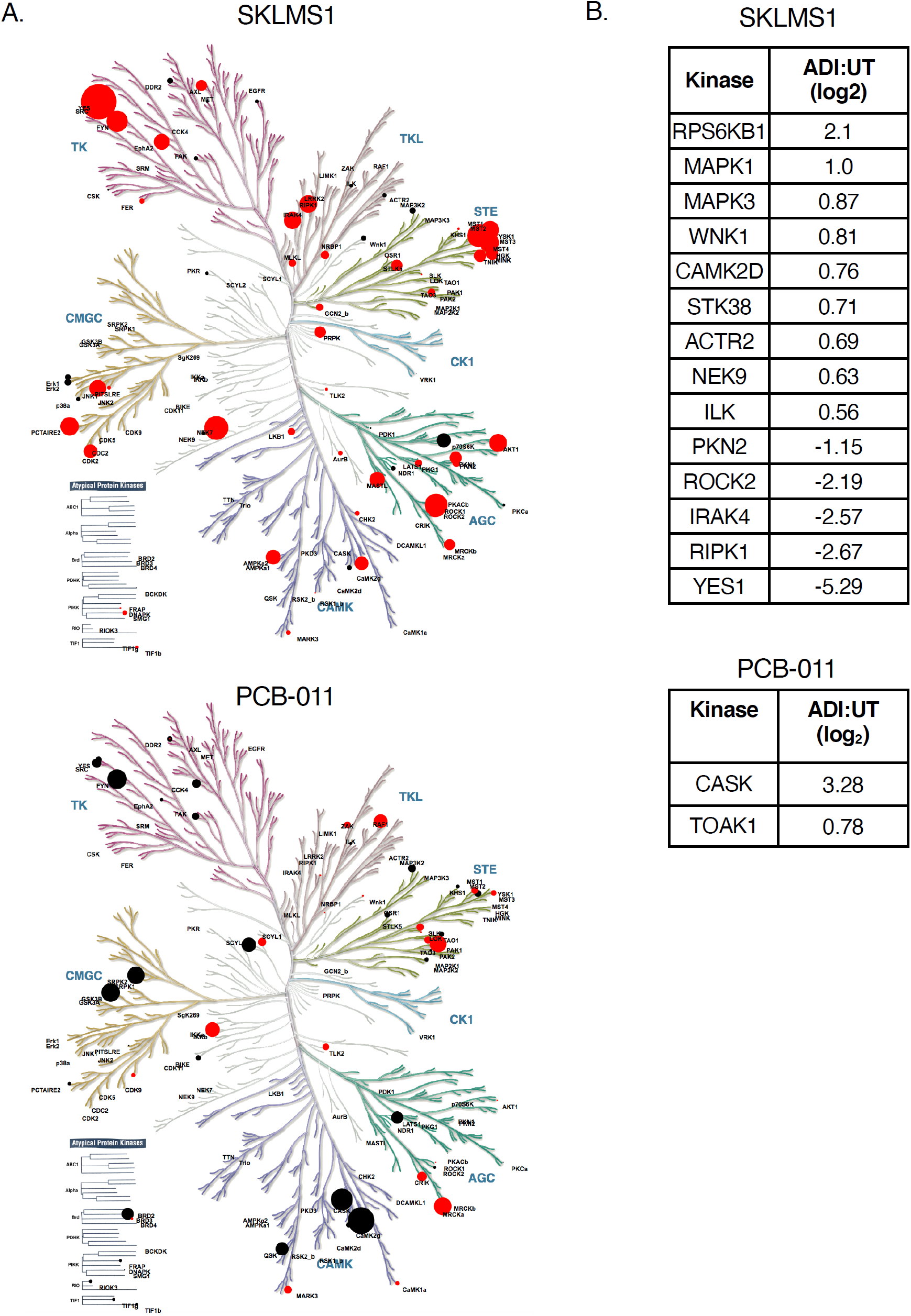
The adaptive kinome of SKLMS1 and PCB-011 cells in response to ADI-PEG20-PEG20 includes ERK. **A)** Kinases identified by ABPP are labeled. Size of the circle indicates relative changes in binding upon ADI-PEG20 treatment and is scaled from the log2 ratio. Color choices are assigned according to the directionality of deviation from no change (black, increased binding; red, decreased binding). Kinome tree illustration reproduced courtesy of Cell Signaling Technology, Inc. (http://www.cellsignaling.com). **B)** Kinases detected by ABPP with a log_2_ fold change >|0.5|.

In order to build a more complete picture of the regulatory networks involved in the response to SKLMS1 and PCB-011 cells to arginine starvation, we performed X2K analysis on the ABPP data (Chen *et al*, 2012). X2K incorporates kinomic and other changes in protein expression to infer regulatory networks. As X2K requires differentially expressed genes as input, all upregulated proteins with a nominally significant *p*-value were included in the X2K analysis of each cell line. Myc, and its activating heterodimeric partner Max, were the two most overrepresented transcription factors in SKLMS1 cells upon ADI-PEG20 treatment, but were much less enriched in PCB-011 cells (Fig. 6A). Myc is stabilized in SKLMS1 cells upon ADI-PEG20 treatment and the cMyc-Max heterodimerization inhibitor 10058-F4 blocks ADI-driven resistance consistent with this regulatory model (Prudner *et al*, 2019b). Network analysis indicated that Myc and Max are driven by upstream activation of ERK1/2 (MAPK1/3, Fig. 6B). ERK can phosphorylate and stabilize Myc (Sears *et al*, 2000; Long *et al*, 2013), supporting this regulatory model upon arginine starvation in SKLMS1 cells. Taken together, the ABPP profiling indicates that adaptive changes in ERK1/2 upon ADI-PEG20 treatment stimulate a cMyc-Max transcriptional network in ADI-PEG20-sensitive SKLMS1 cells, but not in ADI-PEG20-resistant PCB-011 cells.

**Figure 6.**
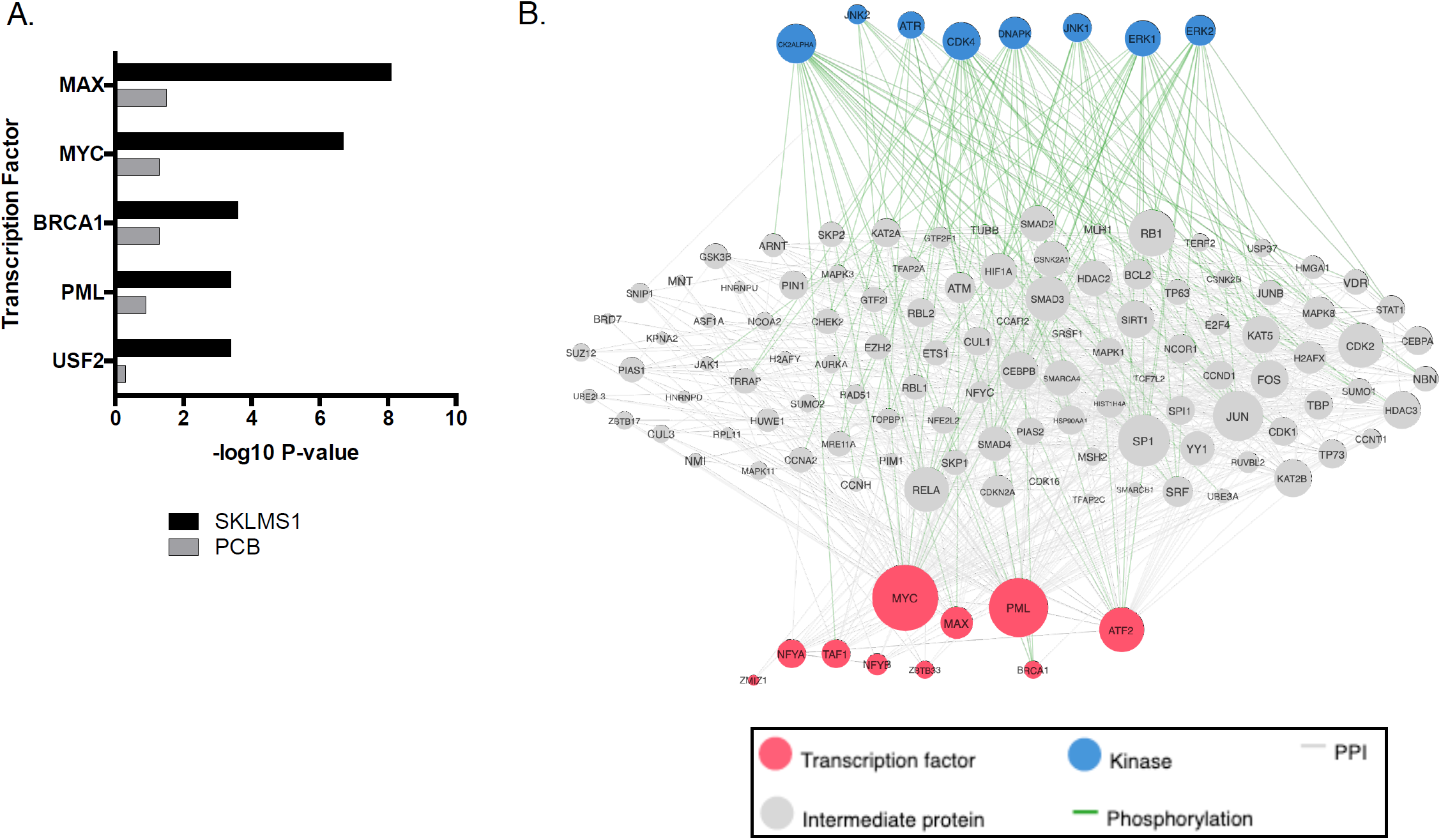
ADI treatment activates a signaling network driving Myc-Max activation. **A)** X2K transcription factor enrichment analysis of SKLMS1 and PCB-011 cells upon ADI treatment. **B)** X2K kinase network analysis based on ABPP profiling of SKLMS1 cells upon ADI treatment.

## Discussion

Acquired resistance to anticancer therapy remains a major challenge and often occurs in the absence of genetic mutations. Many cancers are arginine auxotrophic due to silencing of ASS1 and/or argininosuccinate lyase (Keshet *et al*, 2018). These tumors are sensitive to ADI-PEG20, which converts arginine to citrulline (Przystal *et al*, 2018), but ultimately tumors develop resistance through re-expression of ASS1, metabolic reprogramming (Kremer *et al*, 2017a), and myc stabilization (Prudner *et al*, 2019b). In order to more broadly understand how cells develop resistance to arginine starvation, we focused on identifying the proteomic adaptations that facilitate metabolic reprogramming, and ultimately escape, in ADI-PEG20 sensitive cells,

In this study we employed phosphoproteomics and ABPP to characterize how the proteome of SKLMS1 cells adapts upon arginine starvation. These methods proved to be complementary, and by integrating these data with existing metabolomics data (Kremer *et al*, 2017a), we were able to gain unique insights into the regulatory networks involved in ADI-PEG20 resistance in ASS1-negative sarcoma. In line with existing literature, we identified adaptive kinomic changes in ADI-PEG20-sensitive SKLMS1 cells, including upregulation of ERK1 and ERK2, which stimulate a Myc-Max transcriptional network (Fig. 4A and 6A). Myc in this context has been demonstrated to promote expression of ASS1 (Long *et al*, 2013). Additionally, Myc is able to promote glutamine anapleurosis (Gao *et al*, 2009), but in the setting of arginine deprevation it is not known how proteomic changes facilitate this metabolic reprogramming. We find that regulation of multiple proteins contributes to increased flux from glutamine to OAA (glutamine anapleurosis), direct oxaloacetate production from pyruvate by increasing pyruvate carboxylase (pyruvate anaplerosis), and upregulation of citrate synthase combined with inhibition of lipid synthesis to recycle citrate for TCA anaplerosis (Fig. 4B and 4C).

The reprogramming of cancer metabolism is a critical factor promoting tumorigenesis and drug resistance. Proteomic regulation is essential to metabolic reprogramming (Zhou *et al*, 2012), such as PKM2 tetramerization to promote the Warburg effect (Benjamin *et al*, 2012). While the limited number of cellular metabolites (Mahieu & Patti, 2017) make metabolomic profiling relatively routine, the vast complexity of proteomic regulation remains challenging and the myriad potential mechanisms by which proteomic changes drive metabolic reprogramming remain incompletely understood. Systems level analysis pairing global, unbiased, and integrative proteomic and metabolic analyses have recently been performed in models of plants (Wienkoop *et al*, 2008; Faddetta *et al*, 2018; Blachowicz *et al*, 2019), parasites, and antibiotic resistance (Akpunarlieva *et al*, 2017; Park *et al*, 2016). However, utilization of integrative proteomic-metabolomic analysis has been limited with respect to modeling drug resistance and metabolism in cancer models (Feng *et al*, 2018; Blum *et al*, 2018; A.L. *et al*, 2015; Cai *et al*, 2010). Our exploration of ADI-PEG20 resistance elucidates numerous changes consistent with metabolomics, and finds that these two -omics approaches provide complementary insight.

Importantly, 14% of the proteins detected by ABPP are annotated in UniProtKB as ATP binding, which is an underestimation and is a percentage on par with another comprehensive investigation of ATP binding proteins in *E. coli* (Piazza *et al*, 2018b). ATP is a broad allosteric modulator (Lu *et al*, 2014) and thus the majority of the proteins detected are likely direct, but not enzymatically regulated, ATP interactors. However, protein-protein interactions, which are another indicator of altered protein regulation, can also contribute to the binding and detection by ABPP when using an ATP resin. The imprecision of ABPP is therefore akin to commonly used kinase phosphorylation analyses, which is indicative of regulatory events, but only modestly correlated with kinase activity for even well-studied kinases such as AKT (Ranganathan *et al*, 2007; Tsurutani *et al*, 2006; Vincent *et al*, 2011). Therefore, the sole use of ABPP for the elucidation of complex metabolic networks is unlikely to provide a comprehensive pathway analysis. Combining ABPP with other -omics allows for a more complete understanding of these interconnected systems.

For arginine starvation to become a mainstay of cancer treatment, a full understanding of the metabolic and proteomic adaptation to ADI-PEG20 is needed. The metabolic changes that promote escape from arginine starvation induce a new transcriptional profile as well as significant alterations in protein regulation and activity. These changes that facilitate escape from arginine deprivation also limit the metabolic flexibility of cells. It has yet to be determined if these metabolic adaptations are permanent choices, but as long as the stress of extracellular arginine starvation is present, the ability of a tumor to return to its metabolism of choice is limited. Understanding the metabolic changes that occur upon treatment with metabolically targeted compounds in concert with associated proteomic changes provides insight into cellular resistance mechanisms and may inform the development of more efficient multiagent therapies.

## Materials and Methods

### Materials

All materials were from Sigma unless otherwise noted. ActivX desthiobiotin ATP kinase enrichment kit and BCA assay were from Pierce. Mass spec grade trypsin was from Promega. Amicon Ultra Centrifugal Filters and C18 ziptips were from Millipore. Bondbreaker TCEP, 5mL 7K MWCO Zeba spin desalting columnsm, and formic acid were from ThermoFisher. Oasis HLB 1 cc extraction columns were from Waters. Lysis buffer for phosphorylation analysis was from Cell Signaling Technologies. Sequencing grade trypsin was from Promega. Ni-NTA agarose beads were from Qiagen. We obtained PCB-011 from Charles Keller, PhD. SKLMS1 cells were from ATCC.

### Cell culture

SKLMS1 and PCB-011 were cultured at 37°C in 5% CO_2_ in Minimum Essential Media (MEM) (ThermoFisher Scientific 11095-072) supplemented with 10% FBS (R&D Systems S111560), Penicillin-Streptomycin (1:000) (ThermoFisher Scientific 15140122), and plasmosin (InvivoGen ant-app). Cells were confirmed to be mycoplasma negative with the mycoalert kit (Lonza LT07-418).

### Drug Treatment and Proliferation Assays

SKLMS1 and PCB-011 were seeded at 2,500 cells per well in a 96-well plate one day prior to the assay. Phenol red free media containing 10% FBS was pretreated with 1μg/mL of ADI-PEG20 on day prior to the assay. On the day of the assay, the media was exchanged for ADI-PEG20 pretreated media or fresh phenol free media in the untreated control. Cell proliferation was measured over the course of 72hr in the IncuCyte Live Cell Imaging System and data was analyzed using IncuCyte S3 imaging software (Sartorius Ann Arbor, MI).

### Immuno Assays

For analysis of ASS1 expression cells were seeded at 200,000 cells per well in a 6 well plate and MEM was pretreated with ADI-PEG20 (1μg/mL) one day prior to the beginning of the assay. On the day of the assay the media was exchanged for ADI-PEG20 pretreated media or fresh MEM in the untreated control. After 72hr of treatment cells were lysed with 1x Cell Lysis Buffer (9803, Cell Signaling Technology) according to the reagent protocol. Lysates were run on a ProteinSimple Wes automated western blot using the instrument default settings and the ProteinSimple standard protocol. Protein Simple Compass was utilized for the data analysis. Anti-ASS1(Polaris) antibody was used at 1:1000.

### Activity-based proteomic profiling (ABPP) using ATP resin

Cells in a 10 cm dish were lysed and assayed using the Pierce kinase enrichment kit with the ActivX desthiobiotin-ATP probe per manufacturer’s instructions. Briefly, cells were treated with ADI-PEG20 for 72 hr, trypsinized and pelleted at 1000 x g for 5 min. The pellet was washed once with 5 mL PBS and lysed in 1mL Pierce IP lysis buffer with the included protease/phosphatase inhibitors added. Lysates were desalted with 5mL 7K MWCO Zeba according to manufacturer’s instructions, diluted to 2 mg/mL in lysis buffer, and labeled with 20 uM desthiobiotin-ATP for 10 min at room temperature. Proteins were enriched with the IP lysis buffer plus 8 M urea with 50 uL 50% slurry of the high capacity streptavidin sepharose included in the kit, rotating end-over-end for 1 hour. Resin was washed 3 times with lysis buffer supplemented with 4M urea prior to elution with 500 uL 0.5% SDS, 1% BME in 0.1M Tris pH 6.8 and heated at 95°C for 5 min. A second, identical elution was performed and combined with the first. Lysates were reduced (10 mM DTT, 25 min, 60°C), alkylated with iodoacetamide (18 mM, 30 min) and concentrated with Millipore Amicon Ultra spin columns (UltraCel, 10K MWCO). Proteins were precipitated with 5 volumes MeOH:chloroform (4:1, v/v). The interphase was isolated, washed with MeOH, and proteolyzed with 1.25 ug trypsin in 78 uL of 2% acetonitrile overnight at 37°C at 900 rpm. Samples were acidified with 0.5% formic acid, desalted with C18 zip tips (0.6 uL resin), eluted with 80% acetonitrile, 0.5% formic acid prior to vacuum concentrated to near dryness prior to LC-MS analysis.

### IMAC enrichment of phosphopeptides

Cells in 10 cm dishes were treated with or without ADI-PEG20 (10uM) for 72 h, washed with PBS and lysed in 1X Cell Signaling Cell Lysis Buffer plus 1 mM PMSF. Lysates were sonicated in a water bath on ice for 15 sec and insoluble material was removed with a 14,000 x g centrifugation for 10 min at 4°C. Five 200 ug aliquots of lysate were made for each sample. Each aliquot was desalted with 600 uL GE 2-D Clean-Up kit and processed through to trypsin digest as in ABPP. Lysates were desalted using Oasis HLB columns per manufacturer’s instructions. Samples were then diluted with 1 mL 90% ACN, and phosphopeptides were enriched with 20 uL of Qiagen Ni-NTA slurry for 30 min at 25°C with end-over-end rotation. Beads were washed four times with 1 mL 80% ACN, 0.1% TFA and eluted with 250uL 50% ACN, 2.5% ammonia, and 2 mM phosphate buffer pH 10. Lysates were acidified to pH <3 with formic acid, vacuum concentrated to dryness, desalted with a C18 ziptip using manufacturer’s instructions, vacuum concentrated to dryness, resuspended in 0.5 formic acid and analyzed by LC-MS.

### LC-MS

Samples were analyzed by reverse-phase HPLC-ESI-MS/MS using a nano-LC 2D HPLC system (Eksigent) which was directly connected to a quadrupole time-of-flight (QqTOF) TripleTOF 5600 mass spectrometer (AB SCIEX) in direct injection mode. 3 μL of analyte was loaded onto 3 μl sample loop. After injection, peptide mixtures were transferred onto a self-packed (ReproSil-Pur C18-AQ, 3μm,Dr. Maisch) nanocapillary HPLC column (75 μm I.D. x 22 cm column) and eluted at a flow rate of 250 nL/min using the following gradient: 2% solvent B in A (from 0-7 min), 2-5% solvent B in A (from 7.1 min), 5-30% solvent B in A (from 7.1-130 min), 30-80% solvent B in A (from 130-145 min), isocratic 80% solvent B in A (from 145-149 min) and 80-2% solvent B in A (from 149-150 min), with a total runtime of 180 min including mobile phase equilibration. Solvents were prepared as follows: mobile phase A: 0.1% formic acid (v/v) in water, and mobile phase B: 0.1% formic acid (v/v) in acetonitrile. Mass spectra were recorded in positive-ion mode. After acquisition of ∼1-3 samples, TOF MS spectra and TOF MS/MS spectra were automatically calibrated during dynamic LC-MS & MS/MS autocalibration acquisitions injecting beta-galactosidase (AB SCIEX). Automatic re-calibration of TOF-MS and TOF-MS/MS scans (after every 1-3 samples) guaranteed reliably high mass accuracy and MS detector precision over an extended period of time (weeks) without interruption of the HPLC-MS/MS acquisitions. Two different mass spectrometric acquisition workflows were performed in this study: *1) Data dependent acquisitions (DDA):* for collision induced dissociation tandem mass spectrometry (CID-MS/MS), the mass window for precursor ion selection of the quadrupole mass analyzer was set to ± 0.7 *m/z*. MS1 scans ranged from 380-1250 *m/z* at a resolution of 30,000 with an accumulation time of 250 ms. The precursor ions were fragmented in a collision cell using nitrogen as the collision gas. Advanced information dependent acquisition was used for MS2 (MS/MS) collection on the TripleTOF 5600 at a resolution of 15,000 to obtain MS2 spectra for the 50 most abundant parent ions following each survey MS1 scan. Dynamic exclusion features were based on value M not *m/z* and were set to an exclusion mass width 50 mDa and an exclusion duration of 30 sec. MS2 scans ranged from 100-1500 *m/z* with an accumulation time of 50 msec. *2) Data independent MS2 acquisitions (DIA):* In the ‘SWATH’ DIA MS2 acquisition, instead of the Q1 quadrupole transmitting a narrow mass range through to the collision cell, a wider window of ∼10 *m/z* is passed in incremental steps over the full mass range (*m/z* 400-1250 with 85 SWATH segments, 63 msec accumulation time each, yielding a cycle time of 5.5 sec which includes one MS1 scan with 50 msec accumulation time). SWATH MS2 produces complex MS/MS spectra that are a composite of all the analytes within each selected Q1 *m/z* window. The RAW and processed data associated with this manuscript have been deposited to the ProteomeXchange repository with the identifier PXD017043.<u>Protein identification and MS1 quantification with MaxQuant:</u> Mass spectral data sets were analyzed and searched with MaxQuant (ver.1.5.2) (Cox & Mann, 2008) against the Uniprot Human Reference Proteome. The MS/MS spectra were searched with fixed modification of carbamidomethyl cysteine, variable modifications of oxidation (M), acetylation (protein N-term), and Gln- > pyro-Glu. Phosphorylation (STY) modification was included for the phospho-enriched peptides. Search parameters were set to an initial precursor ion tolerance of 0.07 Da and main peptide search tolerance of 0.0006 Da. Strict tryptic specificity was required with a maximum of two missed cleavages. The minimum required peptide length was set to seven amino acids. Peptide identification FDR was set at 1%. PTM site assignment as well as label-free protein and peptide quantification was performed in MaxQuant. Common proteins contaminants included the CRAPome (Mellacheruvu *et al*, 2013) were removed prior to pathway analysis. The RAW and processed data associated with this manuscript have been deposited to the ProteomeXchange repository with the identifier PXD017043.

### Skyline Data Analysis

Skyline software (https://skyline.ms/project/home/begin.view?) was used to manually examine and quantify DIA data. Spectral libraries were generated in Skyline using the DDA database searches of the raw data files. Raw files were directly imported into Skyline in their native file format and only cysteine containing peptides were quantified.

### Gene set enrichment analysis (GSEA)

GSEA (Subramanian *et al*, 2005) was performed using GSEA version 2.2.2 from the Broad Institute at MIT. Parameters used for the analysis were as follows. Datasets with protein expression fold changes due to drug treatment were testing for enrichment against BioCarta, Hallmark, Reactome and KEGG gene sets. Number of permutations was set to 1000 to calculate *p*-value and permutation type was set to gene_set. All basic and advanced fields were set to default. Phosphopeptides were assigned to specific genes based on the MaxQuant annotation.

### Kinome tree plot

KinMap (Eid *et al*, 2017) was used to generate the kinome tree based on relative expression in the ABPP dataset.

### X2K

X2K (Chen *et al*, 2012) was performed using default parameters and Networkin as the kinome database. Input was all proteins upregulated by ADI-PEG20 (nominal p-value ≤0.05) in SKLMS1 cells in the ABPP dataset.

## Acknowledgements

BAVT, CB, and RR were supported by NCI RO1 CA227115, as well as research grants from CJ’s Journey, The Sarcoma Foundation of America, and The Sarcoma Alliance for Research and Collaboration and The Josephine Norcia Riley Angiosarcoma Awareness Inc. Grant. JH acknowledges funding and support from AB SCIEX.

## Abbreviations

αKG: alphaketoglutarate
ABPP: Activity-based proteomic profiling
AcCoA: Acetyl-CoA
ACN: acetonitrile
ADI-PEG20: Arginine diminase
PEG-2000 Asp: aspartate
ATP: adenosine triphosphate
BME: B-mercaptoethanol
Cit: citrate
CS: citrate synthase
DDA: Data-dependent acquisition mass spectrometry
DIA-MS: Data-independent acquisition mass spectrometry, aka SWATH
DTT: dithiothreitol
FASN: fatty acid synthase
FDR: False discovery rate
GBM: glioblastoma multiforme
IsoCit: isocitrate
IDH: isocitrate dehydrogenase
Mal: malate
MAPK: mitogen-activated protein kinase
MDH: malate dehydrogenase
Mito: mitochondria
Myc: Myc-C
OAA: oxaloacetate
PC: pyruvate carboxylase
PDHA: pyruvate dehydrogenase, Alpha-1
PMSF: phenylmethylsulfonyl fluoride
TFA: trifluoroacetic acid
Veh: vehicle

